# Observations of the Effect of Scopolamine on Hippocampal CA1 Place Cell and Network Properties in Freely Moving Mice Using Miniscope Imaging

**DOI:** 10.1101/2020.05.27.120352

**Authors:** Dechuan Sun, Ranjith Rajasekharan Unnithan, Chris French

**Affiliations:** Department of Medicine, The University of Melbourne; Department of Electrical and Electronic Engineering, The University of Melbourne

## Abstract

The hippocampus and associated cholinergic inputs regulate spatial memory in rodents. Muscarinic blockade with scopolamine results in cognition deficits usually attributed to impaired memory encoding, but effects on memory retrieval are controversial. Here, we simultaneously recorded hundreds of neurons in mouse hippocampal CA1 using calcium imaging with a miniatured fluorescent microscope to study place cell and ensemble neuronal properties in a linear track environment. We found decoding accuracy and ensemble stability were significantly reduced after the administration of scopolamine. Several other parameters including the ***Ca***^**2+**^ event rate, number of total cells and place cells observed, spatial information content were affected including a small increase in running speed. This study enhances the understanding of cholinergic blockade on spatial memory impairment.

## 1. Introduction

In rodents, memory formation and retrieval critically rely on the hippocampus (HIP) (Buzsáki et al., 1990; Izquierdo et al., 1997; Wiltgen et al., 2010; Carr et al., 2011). Place cells are hippocampal pyramidal neurons that activate in response to position and have been shown to have an important functional role in encoding and decoding spatial position (Dragoi et al., 2006; Pfeiffer et al., 2013; Wikenheiser et al., 2015). Cholinergic receptors are abundantly expressed in the brain, and the modulation of spatial memory in rodent hippocampus is dependent on cholinergic inputs, especially muscarinic acetylcholine receptors (mAChRs) (Riekkinen et al., 1997; Brazhnik et al., 2003; Svoboda et al., 2017). Scopolamine blockade of mAChRs, has been found to greatly impair memory encoding, but its effect on memory retrieval is less clear (Svoboda et al., 2017). Some studies show no or very little influence of scopolamine on memory decoding (Riekkinen et al., 1997; Deiana et al., 2011; More et al., 2016), while Huang et al. (2011) report effects on both encoding and decoding. Additionally, the mechanism of the effect of scopolamine on memory retrieval is poorly understood. Intracranial electrodes arrays are commonly used to record local field potentials and single unit activity and allow detailed observations of cognition-related processes in terms of neural firing patterns, but the number of cells and spatial position in the brain is generally limited. We used *in vivo* calcium imaging to record the activity of neural ensembles in the hippocampal CA1 region of freely running mice with the miniscopes (Ghosh et al., 2011; Aharoni et al., 2019), which allowed the simultaneous recording of ***Ca*^2+^** activity of a large population of neurons. We demonstrated that the blockade of mAChRs severely impaired the decoding accuracy with an animal running through a linear track, attributable to impaired stability of the hippocampal neural ensemble. Several parameters including ***Ca*^2+^** rate, spatial information, total neuron number as well as the animal’s running speed were also affected.

## 2. Materials and Methods

All surgical and experimental procedures were approved by the Florey Animal Ethics Committee (No. 18-008UM) and were conducted in strict accordance with Australian Animal Welfare Committee guidelines.

### 2.1 Subjects

Five naive adult male C57BL/6 mice aged 12 weeks were obtained from WEHI (Melbourne, VIC) and housed in the Biological Research Facility of the Department of Medicine, Royal Melbourne Hospital, University of Melbourne. All animals weighed 24-25g at the time of surgery and then were housed individually. The facility was maintained on a 12-12h light-dark schedule (lights on: 7:30 am to 19:30 pm) with water and standard mice chow *ad libitum*. All procedures were conducted in the daytime.

### 2.2 Drugs Administration

Scopolamine hydrobromide (Sigma-Aldrich, USA) was dissolved in 0.9% saline and injected intraperitoneally at a volume of 1 mg/kg (Newman et al., 2017).

### 2.3 Stereotaxic Surgery

The surgical procedures had two components – virus infusion and grin lens implantation.

#### Virus infusion

pAAV.Syn.GCaMP6f.WPRE.SV40 virus (titer: 2.2 × 10^13^ GC/mL, obtained from AddGene, USA) was injected into dorsal hippocampus (AP -2.1, ML +2.1, DV -1.7 relative to bregma) through a custom made injecting system over a duration of 15 minutes. The virus was firstly loaded into a 1.5 mm OD, 0.9 mm ID capillary, which was fabricated on a Sutter P-1000 electrode puller to make a sharp tip (diameter: 20-50 um); sealed with silicon oil (Sigma-Aldrich, USA) at the open end; and a brass round rod (diameter: 0.8mm; Albion Alloys, UK), which was fixed on a precise 3D positioner finally fitted into the capillary to control the volume of the virus injected. See Figure 10 for diagram of the injecting system. The virus injector was left in place for additional 10 minutes to allow for viral diffusion. After stitching the wound, the animal was then left for one week to recover and to allow fluorophore expression.

**Figure 1.**
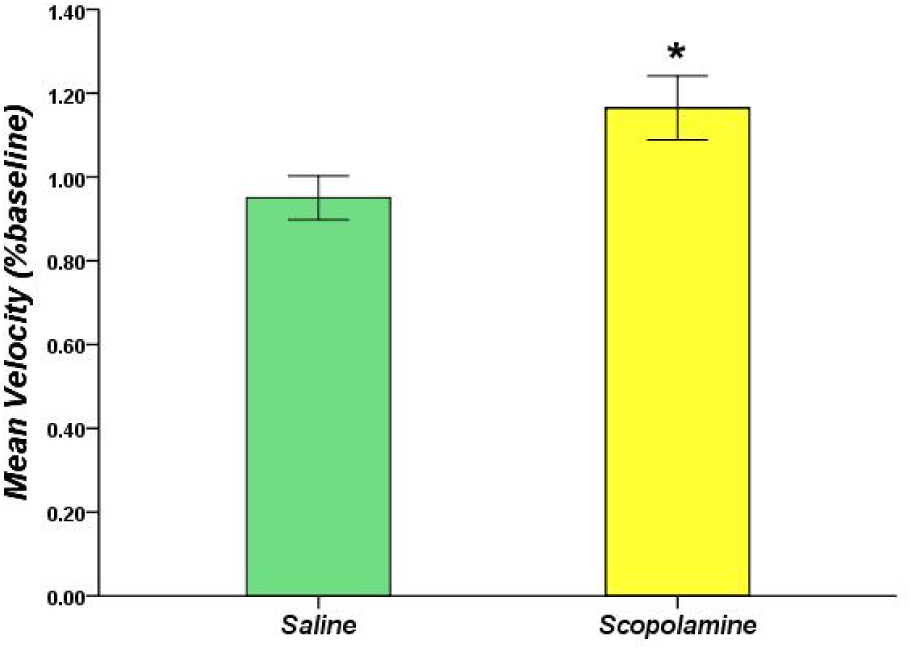
The percentage variation of the animal’s running speed after injecting saline or scopolamine (1 mg/kg) with respect to the baseline self-control. * P<0.05 represents the significant difference between scopolamine and saline groups.

**Figure 2.**
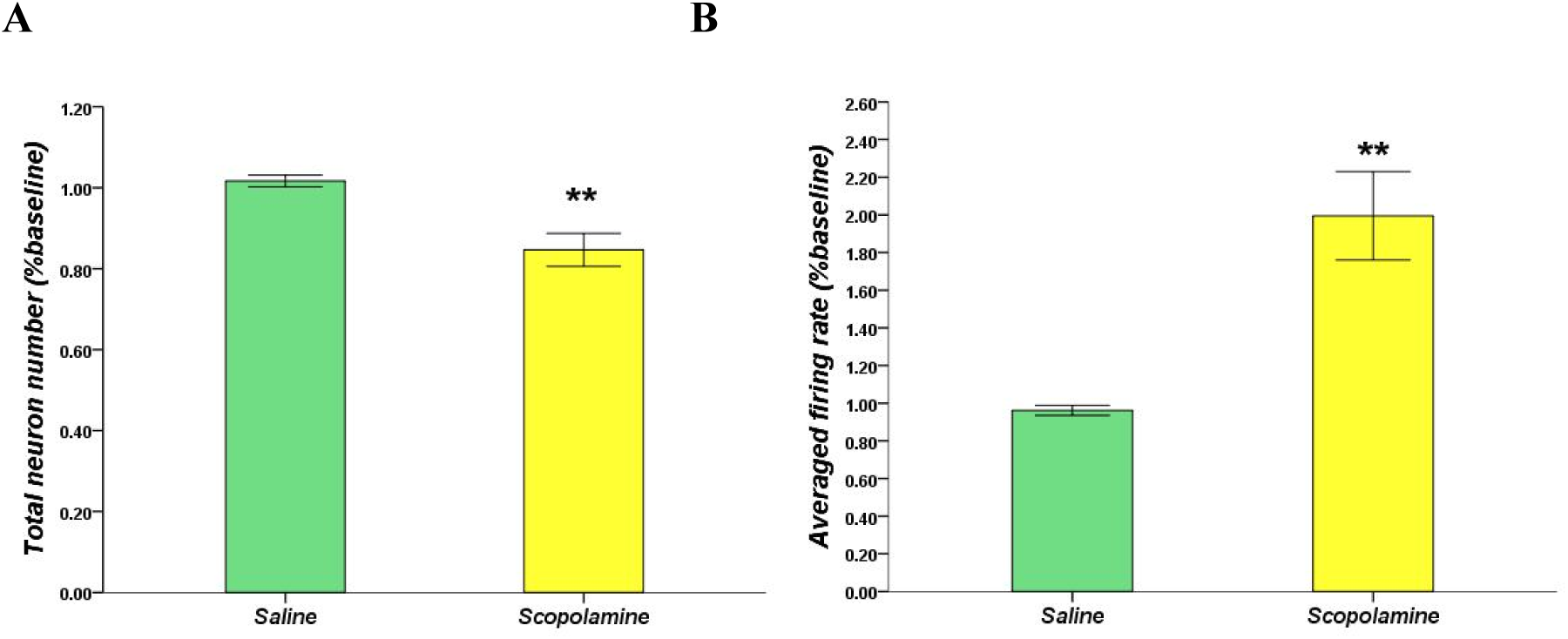
(A)The animal’s total neuron number within the field of view significantly reduced after injecting scopolamine (1 mg/kg) with respect to saline. (B)The scopolamine (1 mg/kg) dramatically enhanced neurons’ averaged *Ca*^2+^ event rate compared with saline. ** P<0.01 represents the significant difference between scopolamine and saline groups.

**Figure 3.**
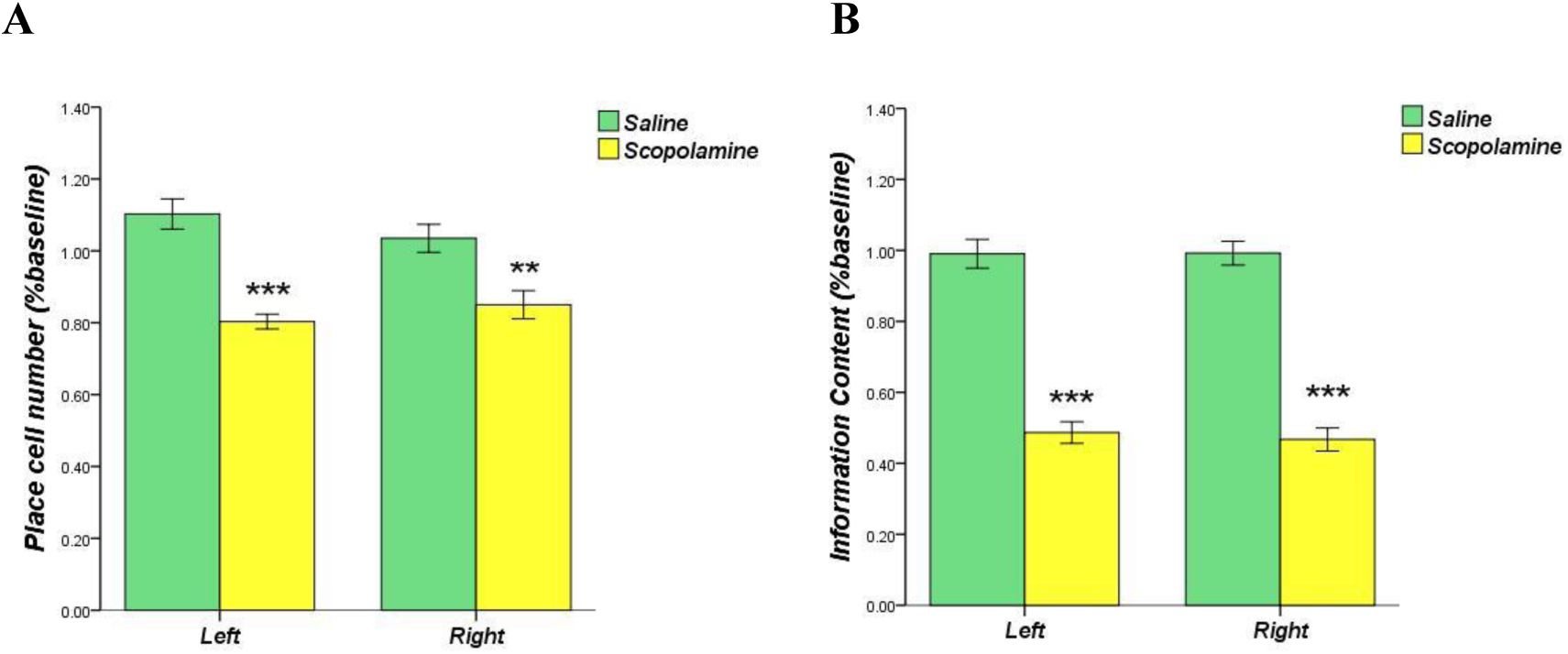
(A) The place cell numbers measured declined significantly after the administration of scopolamine (1 mg/kg) compared with saline during both left-towards and right-towards running directions. (B) The scopolamine (1 mg/kg) significantly depressed the neurons’ spatial information content with respect to saline. ** P<0.01, *** P<0.001 represents the significant difference between scopolamine and saline groups.

**Figure 4.**
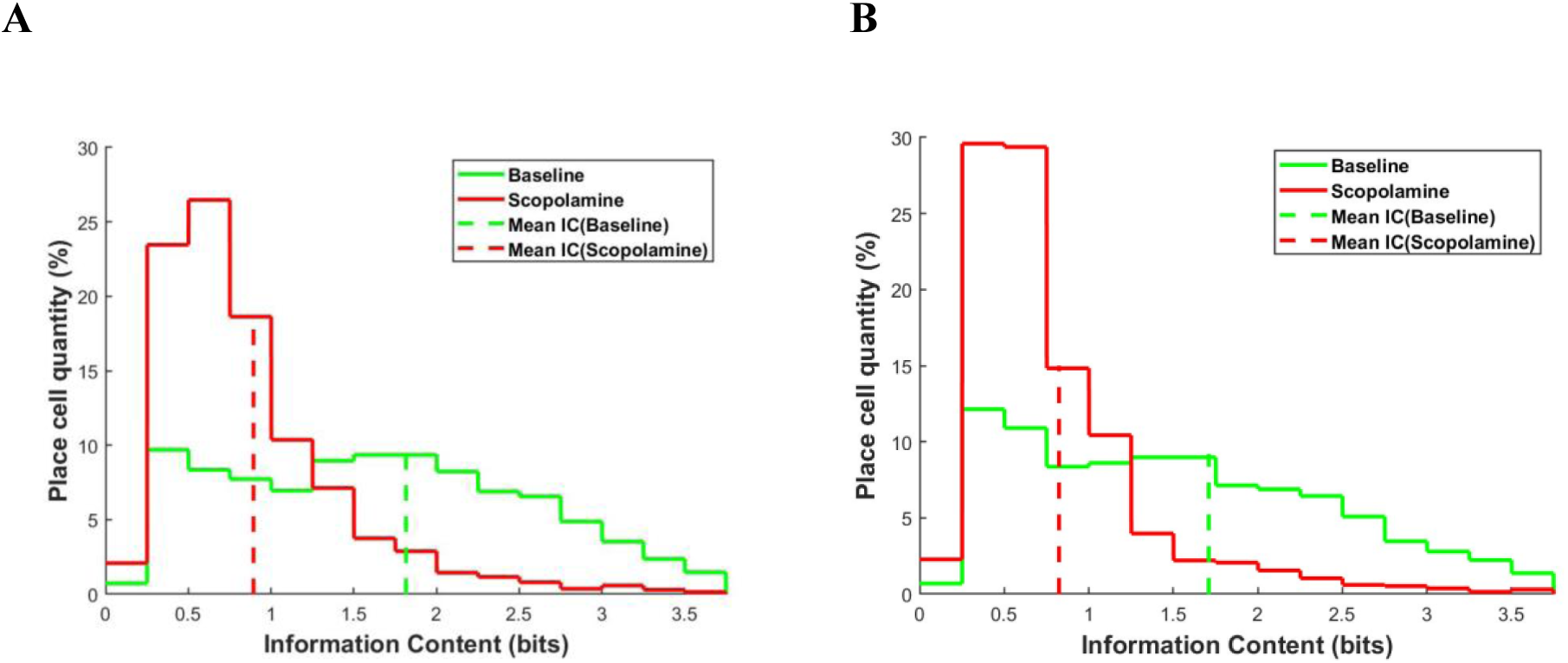
The averaged information content measured across five animals before (green solid line) and after (red solid line) scopolamine (1 mg/kg) injection in left-towards running direction (A) and right-towards running direction (B). The dashed vertical line represented the averaged mean.

**Figure 5.**
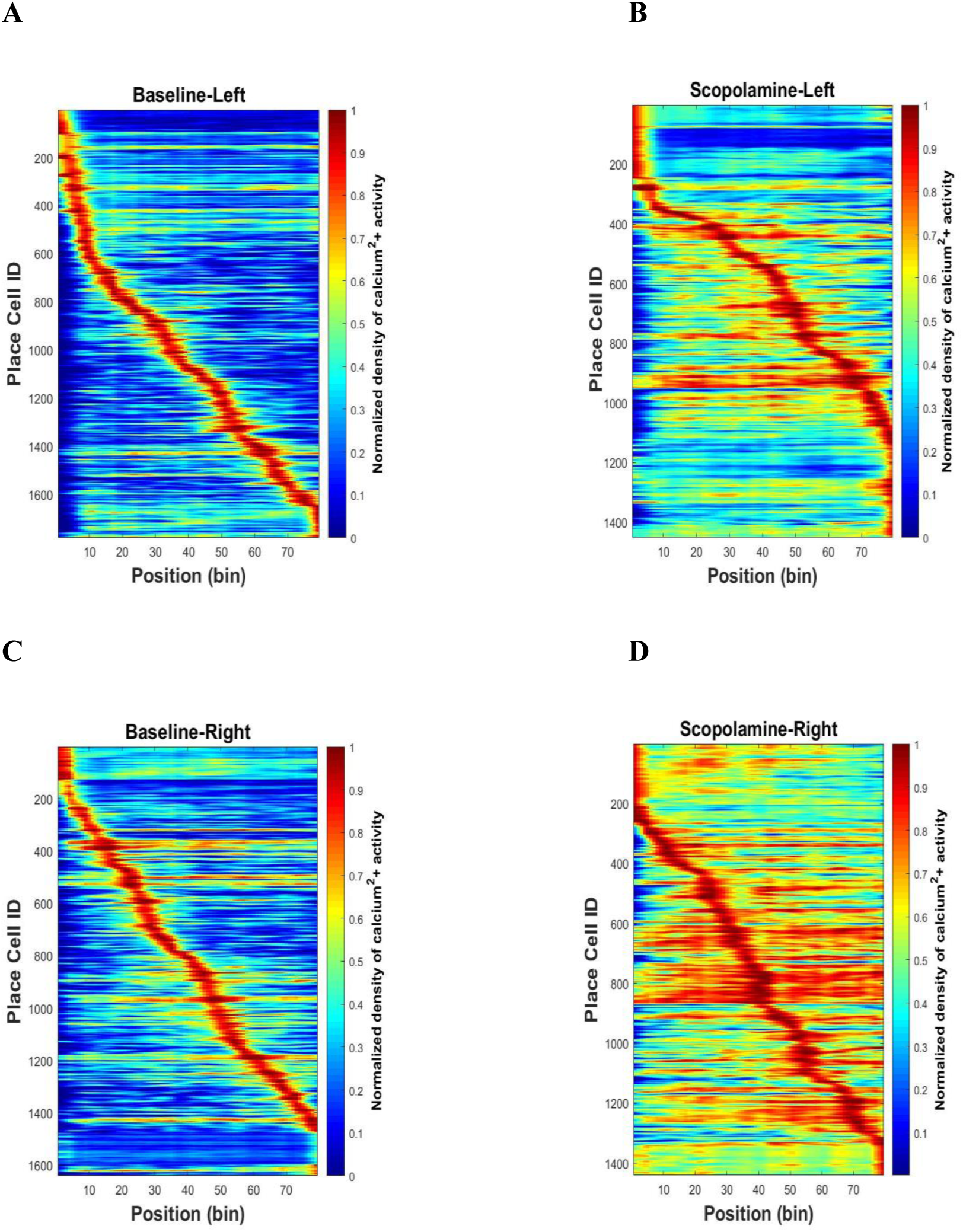
Normalized place field map (See method) when the animal was running on the linear track from right to left before and after scopolamine (1 mg/kg) injection (A, B) and from left to right (C, D). The bin size was 2cm. The location sensitivity of the place cells decreased after drug administration.

**Figure 6.**
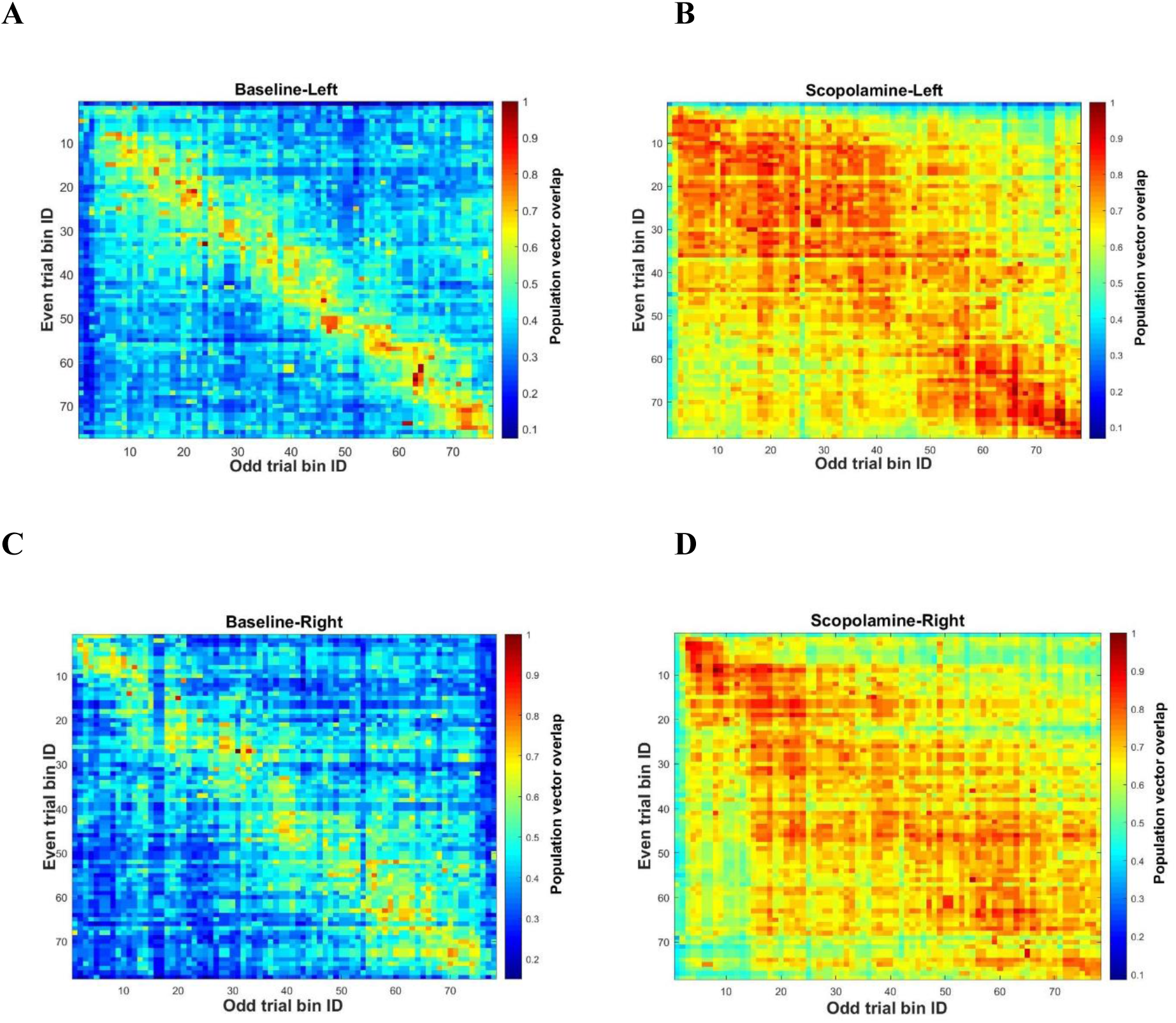
Normalized population vector overlap (PVO) of the place cells between odd trials and even trials before and after the scopolamine (1 mg/kg) administration in left-towards (A, B) and right-towards (C, D) running directions. The PVO increased dramatically after drug injection.

**Figure 7.**
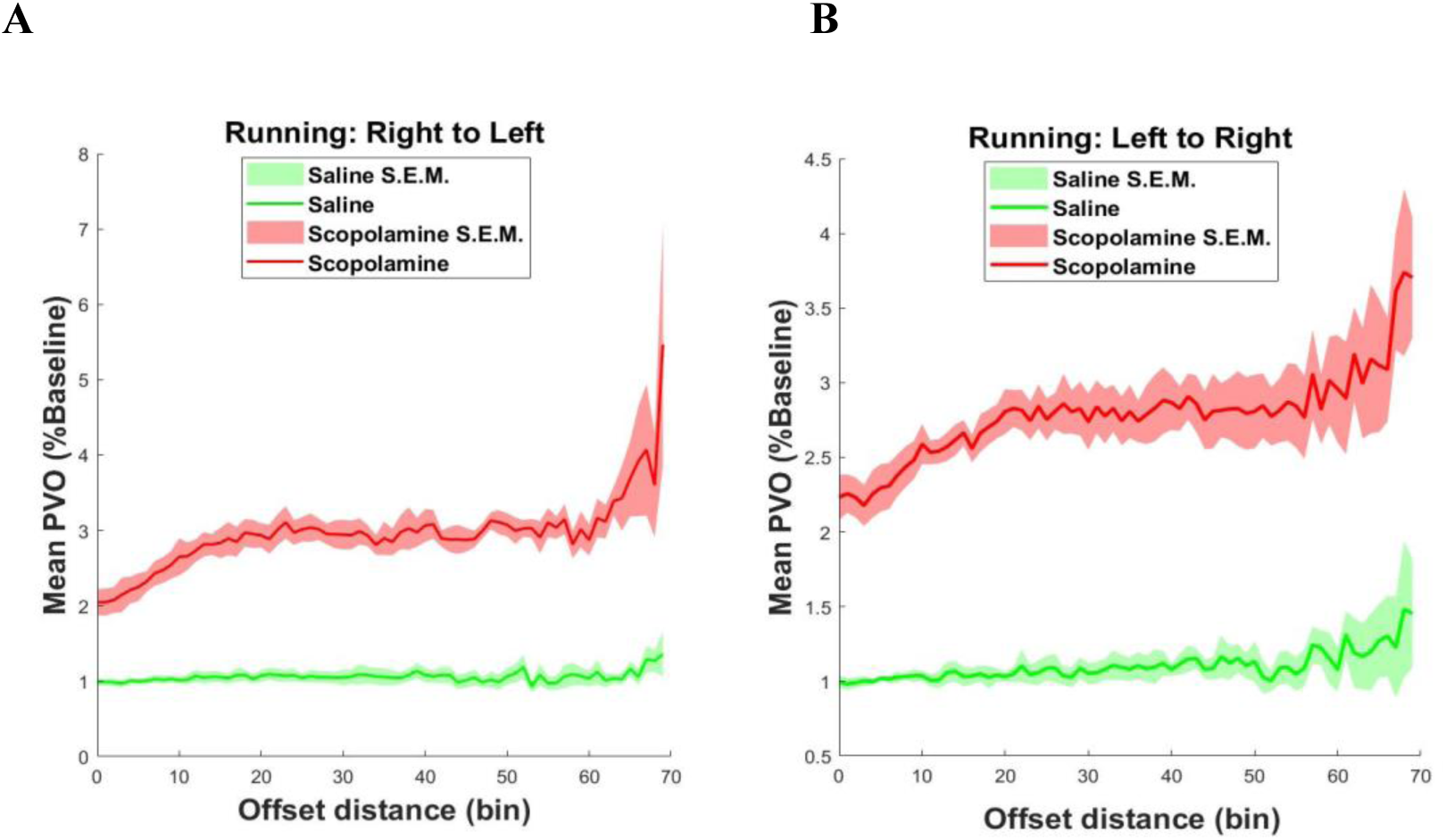
The mean odd-even trails PVO averaged over 5 animals with respect to the distance offset when the animal run from right to left (A) and from left to right (B). After the administration of scopolamine (1 mg/kg), the mean PVO elevated significantly across different distance offset in both running directions (**p<0.01). The shadowed area showed the standard error mean.

**Figure 8.**
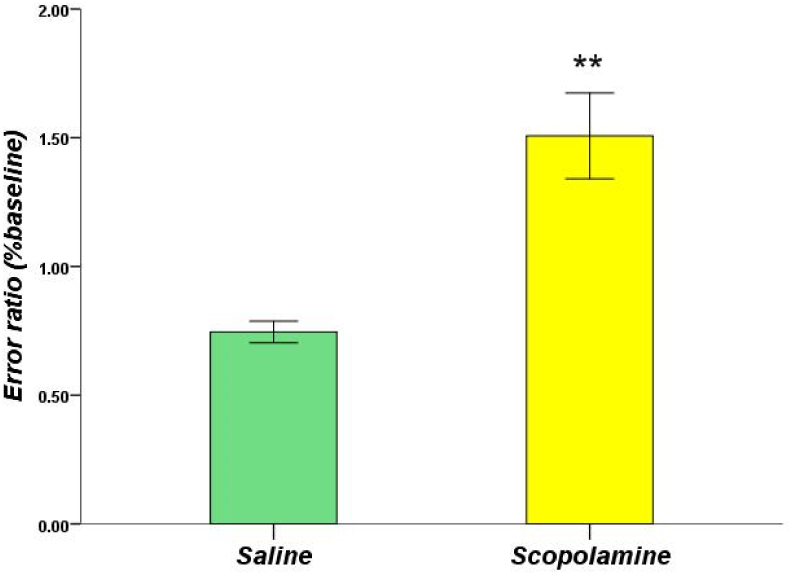
The percentage variation of the error ratio after injecting saline or scopolamine (1 mg/kg) with respect to the baseline self-control. ** P<0.01 represents the significant difference between scopolamine and saline groups.

**Figure 9.**
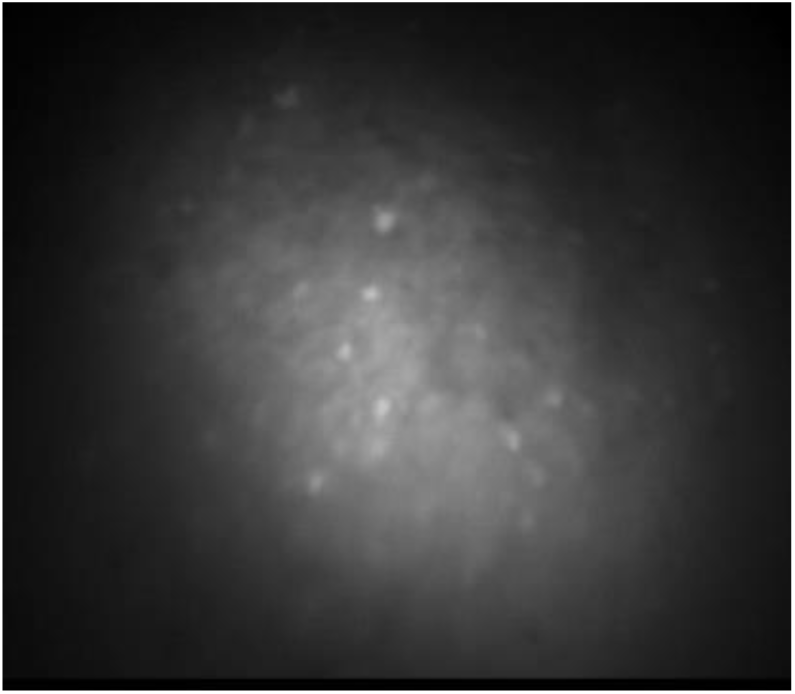
An example of the raw fluorescent data.

**Figure 10.**
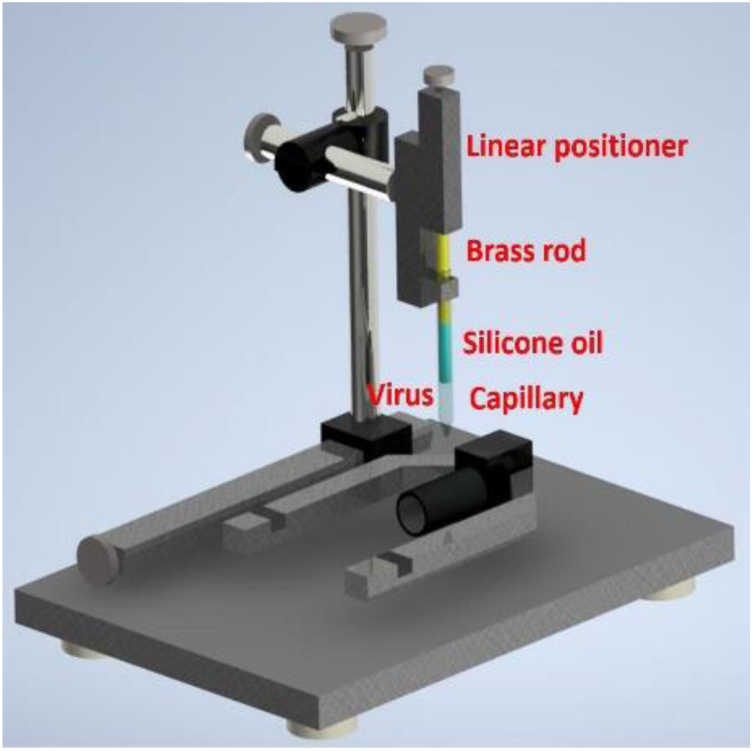
Custom made virus infusion instrument.

#### GRIN lens implantation

Two 1mm screws were implanted (AP +1.8, ML -2.5; AP -2.8, ML -0.8) to serve as anchors. A small window of skull was removed by using a 2mm drill bur, centred at (AP -2.1, ML +1.6) and the exposed dura was cleaned with fine tweezers. A 27-guage blunt needle was used to aspirate the above cortex to expose the vertical striations of the hippocampal fimbria, with artificial cerebrospinal fluid dropped constantly during the procedure to provide a clear operating field. Using the most posterior point of the edge of the drilled hole (next to lambda side) as a reference, the grin lens (0.23pitch, #64-519, Edmund Optics) was implanted 1.35mm deep, touching the surface of exposed tissue. Superglue was applied surrounding the lens to prevent movement and dental cement was built over the glue for support, the lens was then covered with fast setting silicone adhesive (Dragon Skin® Series, USA). After the surgery, the animal was injected with carprofen (5mg/kg) and dexamethasone (0.6mg/kg, Sigma-Aldrich, USA) intraperitoneally everyday, and provided with enrofloxacin water (1:150 dilution, Baytril®, USA) for one week. Four weeks later, a small metal baseplate was mounted on animal’s head to support the miniscope, and the miniscope (focal length of the inside achromatic lens:7.5mm, #45-407, Edmund Optics) was locked in the position at the optimal focal distance.

### 2.4 Animal Training

Before training, the animals were handled approximately 10 minutes twice a day in the daytime and weighed after each handling session for 5 days, to familiarise the animal and the averaged weight could be assumed as a reference of free-feeding weight. A food restriction regimen was implemented to keep the animal at 85% of its original average weight (Kermani et al., 2018). The animal was then trained to run back and forth on a 1.6m linear track with clues painted on the walls for food reward while wearing the miniscope, and a small food pellet was given to the animal once it could rapidly run through the track without wandering. During each training session, the animal performed up to 30 trials within 40 minutes.

### 2.5 Experimental Procedure

All the recordings were performed during daytime. The animal was brought into a silent recording room 30 minutes before the start of recording to become familiar with the surrounding environment. After mounting the miniscope, the animal was put back to its cage for 5 minutes and then moved to the linear track to freely explore the space for 30 minutes. Then the linear track was cleaned with 80% ethanol to eliminate scent clues, and 12 running trials were recorded as a baseline control. Imaging frames were recorded with miniscope acquisition software (UCLA Miniscope, 2017). The excitation LED intensity was set to proper value with a sampling rate of 30 FPS. The animal was then injected with saline or scopolamine, replaced in its cage for 20 minutes and then performed another 12 running trials. A camera fixed overhead was synchronized with miniscope to record the animal’s position and the miniscope cable was suspended over the linear track through a custom-made commutator.

### 2.6 Processing of Calcium Imaging Data

#### Image Pre-processing and Calcium Activity Deconvolution

A non-rigid motion correction algorithm was applied first to implement image registration (Pnevmatikakis et al., 2017). Constrained Non-negative Matrix Factorization for microendoscopic data (CNMF-E) was utilized to identify and extract each neuron’s spatial boundary and calcium activity (Zhou et al., 2018). A fast deconvolution algorithm was then utilized to deconvolve the calcium activity to estimate neural spike-activity (Friedrich et al., 2017). This algorithm sometimes produces low-amplitude “partial spikes”, which were removed by setting a small threshold. This threshold is chosen by observing the distribution of the deconvolved signals across cells (Pennington et al., 2019). We refer to this deconvoluted signal as “temporal spike activity”.

#### Place field map

The position of the animal’s head and running speed were detected by using custom Matlab script. We separately analysed the data of both left-to-right (LR) and right-to-left (RL) running directions. To analyse the neural spatial spike activity, the linear track was divided into several 2cm bins (the bins on each end were discarded), and a speed threshold of 8cm/s was set, then the bins occupancy was calculated as well as the calcium event rate (the number of spikes in each bin) of each neuron in all bins, and a Gaussian smoothing kernel (σ = 1.5 bins, size = 5 bins) was applied. The place field map for each neuron was measured by dividing each neuron’s smoothed spatial spike activity by the smoothed bins occupancy, with the maximum value defined as the place field’s position (Rubin et al., 2015).

#### Spatial information content and place cells

The neuron’s spatial information content was defined as (Markus et al., 1994):

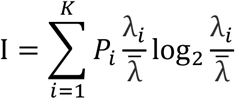

Where K is the number of bins; *P*_*i*_ is the occupancy ratio of the bin i; *λ*_*i*_ is the neuron’s calcium event rate in bin i;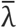 is the mean calcium event 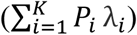.

We used the unsmoothed bins occupancy and the calcium event rate to calculate the spatial information content for each neuron and then shuffled the animal’s position as well as neuron’s temporal spike activity for 800 times. A place cell was defined as the neuron whose spatial information content was above chance (p<0.05) with respect to the shuffling results.

#### Odd & Even trials Population Vector Overlap

To look at the similarity degree of neuron’s firing pattern within one session, we calculated the population vector overlap (PVO) between odd and even trials (Ravassard et al., 2013).

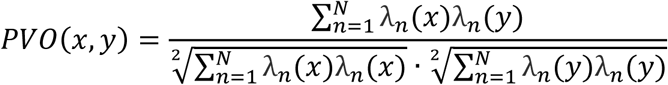

Where N is the number of total neurons; x and y are different bins; *λ* is the place field map.

#### Decoding

A leave-one-out naïve Bayesian decoder was utilized to estimate the position of the animal based on neuronal temporal spike activity (Zhang et al., 1998).

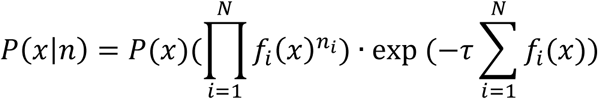

Where n is the current input temporal spike activity; x is the bin number; P(x) is the occupancy ratio of bin x; *f*_*i*_(*x*) is the average calcium event rate of neuron i at bin x; *τ* is the time window length of the input temporal spike activity.

#### Statistics

A two-tailed t-test was performed using SPSS (IBM, Armonk, NY, USA) for all the measures. The level of significance was set at p<0.05. All results were showed as means ± standard error of mean (S.E.M) of the percentage variation from the baseline

## 3. Results

We want to explore how does the antagonists of muscarinic cholinergic transmission affect the activity of the neural ensemble in hippocampal CA1 by observing the fluorescent calcium signal of each neuron within a large field of view. We injected an AAV encoded calcium indicator into hippocampal CA1 of mouse and collected the fluorescent signal as the mouse run back and forth in a linear track before and after the administration of saline and scopolamine. Figure 9 showed an example frame of the raw fluorescent data. After deconvolving the neural spike activity, we compared the firing patterns of the place cell ensembles between control and drug groups, including total cell numbers, place cell numbers, place cells’ total *Ca*^2+^ event rate, place field map, spatial information, odd and even trials population vector overlap, decoding error, as well as the animal’s running speed. All results were showed as a percentage change after injecting saline or scopolamine with respect to the baseline self-control.

### Scopolamine increased running speed

The animal’s average running speed was calculated using all the data on both running directions except the bins on each end of the linear track. The scopolamine treated mice had a slightly higher running velocity (increased to 116.48% with respect to baseline, SEM: 7.62%) compared with saline treated ones (decreased to 95.02% with respect to baseline, SEM: 5.23%) by applying a 2-tailed t-test (p<0.05; Fig1).

### Scopolamine reduced the total cell number of cells detected but increased the *Ca*^2+^ event rate

We first studied the effects of blocking mAChR on total neuron number as well as the average *Ca*^2+^ event rate. Saline injection had no significant effect on neuron number. Scopolamine significantly (2-tailed t-test, p<0.01) reduced the total number of detected neurons to 84.7%, (S.E.M: 9.10%) of the baseline level (Fig 2A). Before injecting scopolamine, there were about 640 neurons (S.E.M:46.0) in each animal and the number decreased to 541 (S.E.M:46.0) after injection. Interestingly, the remaining neurons with calcium signal became more active after the administration of scopolamine, and the average *Ca*^2+^ event rate was greatly enhanced (199.5%, S.E.M: 23.5%) compared with saline (96.2%, S.E.M: 2.66%; 2-tailed t-test, p<0.01; Fig 2B). Slow calcium firing patterns (1.05 HZ, S.E.M:0.16) occurred at baseline but the *Ca*^2+^ event rate almost doubled after scopolamine administration (1.96 HZ, S.E.M:0.10).

### Reduction of place cell number and spatial information content occurred with scopolamine administration

Head direction plays an important role in rodent spatial navigation (Muir et al., 2002), so we separately analysed the data on both left and right running trials. Spatial information content (whether the neuron had one or several specific firing locations or not) was used as the criterion for location specificity. A small increase in place cells was associated with saline injections in both RL (110.25%, S.E.M: 4.21%, P>0.05) and LR running sessions (103.49%, S.E.M: 3.90%, P>0.05; Fig 3A), and probably because the location sensitivity is reinforced with more running trials. However, both ratios declined after scopolamine injection (80.32%, S.E.M: 2.04%, p<0.001 and 84.98%, S.E.M: 3.91%, p<0.01 separately; Fig 3A). During the baseline observation, a mean of 356 place cells (S.E.M: 38) was observed in each animal on the RL running session and 328 place cells (S.E.M: 30) on the LR running session. After the administration of scopolamine, the numbers reduced to 283 (RL, S.E.M:26) and 279 (LR, S.E.M:31) separately. Spatial information content of the place cells was then quantified. Scopolamine greatly impaired location sensitivity, with the spatial information content decreasing by more than half (RL, 48.68%, S.E.M: 6.79%, p<0.001 and LR, 46.72%, S.E.M: 7.28%, p<0.001; Fig 3B). During baseline recording, average information content was 1.81 bits (S.E.M: 0.17) and 1.71 bits (S.E.M: 0.17) on RL and LR running directions, which decreased to 0.88 bits (S.E.M: 0.10) and 0.79 bits (S.E.M: 0.09) after scopolamine administration. Figure 4 showed the average spatial information content before and after drug administration.

### Neuronal Ensemble Stability is Impaired by Scopolamine

To further explore the effects of blocking mAChR on neural ensemble activity, we first calculated the place field map (see Method; Fig.5) when the animal was running through the linear track. Before injecting scopolamine, most of place cells had relatively consistent firing locations in both running directions. Interestingly, although most of the place cells still showed a certain level of place sensitivity after scopolamine, it was greatly impaired compared with the baseline, and the neurons’ “spontaneous” firing became stronger. In order to quantify the level of ensemble stability, we analysed the population vector overlap (PVO) between the odd trials and even trials data within every session, which revealed the overlap of ensemble activity in these two different conditions. In Figure 6, there is a very clear diagonalization feature during the baseline in both running directions, showing a specific and consistent firing pattern in both even and odd trials. However, due to mAChR inhibition, this diagonalization feature disappeared, and the stability of the neural system degenerated. We then plotted the mean PVO ratio with respect to the distance offset (Fig.7). After injecting saline, the PVO ratio changed little compared with the baseline and it was quite stable with very small standard error mean, and once the scopolamine was injected, the PVO ratio increased significantly in both RL (p<0.01) and LR (p<0.01) running directions. Besides, the PVO ratio tended to increase with respect to the offset distance, which may be because of the neural decoding system degeneration.

### Scopolamine decreased decoding accuracy

The stability of the neural ensemble greatly influenced decoding. We analysed the animal’s neural decoding accuracy by using a Bayesian decoder to predict the animal’s position (see Method). In this case, the entire neuron population was considered in the decoding instead of just one neuron and we only used the place cells to train the decoder. The error ratio decreased (25.60%, S.E.M: 4.27%) after saline injection with respect to the baseline and significantly enhanced after dealing with scopolamine (150.68%, S.E.M: 16.68%; p<0.01). Before mAChR was blocked, the average estimation error was 1.3 cm/frame (S.E.M: 0.08), while the error enhanced to 3.16 cm/frame (S.E.M: 0.17) after blocking, which increased almost 2.5 times.

## 4. Discussion

In this study we have observed the effects of scopolamine on calcium signalling in CA1 neurons in mouse hippocampus during free movement in a linear track. Scopolamine had quite striking effects on cellular and ensemble behaviour, disrupting place cell specificity, which we interpret as a disruptive effect on decoding of neural correlates of spatial memory.

Scopolamine has been used frequently previously to study behavioural, neurochemical and electrophysiological effects of disruption of the muscarinic cholinergic system known to be critical for hippocampal function. It has strong amnestic effects and has been used extensively as a model for the memory dysfunction seen in neurodegenerative conditions such as Alzheimer’s disease associated with cholinergic transmission impairment. Previous studies have emphasized impairment of encoding of memory, but in this study we have examined effects on place cells and related ensembles after encoding so it would be expected that the changes in place cell properties we think are best ascribed to impairment of the decoding process, an effect that has been found previously (Huang et al, 2010).

We used times of onset and dosing levels similar to previous studies (Klinkenberg and Blokland, 2010; Falsafi et al., 2012). We found that the animal’s average running speed increased slightly after injecting scopolamine, differing from previous findings (Douchamps et al., 2013; Newman et al., 2017), and this may be due to a higher dose of scopolamine used or the animal studied (mouse rather than rats). Increase in locomotion speed was observed in Bushnell (1987) with dose dependent enhancement in mobility with ∼25% increase with 1mg/kg, but no error estimates are provided. (Bushnell, 1987; As hippocampal CA1 neuronal firing is affected by running speed in place fields (Geisler et al., 2007), we adjusted for this variable by calculating the animal’s running speed distributions of the baseline and after injection, and only analysing frames in which the animal was moving above a designated velocity, to minimise possible effects.

CA1 pyramidal cells display clear location-specific firing (O’Keefe and Dostrovsky, 1971; see Moser et al., 2008 for review), while interneurons appear to have relatively weaker spatial modulation (Hangya et al., 2010). In the current study we were unable to distinguish between these two cell types, but this is unlikely to significantly affect the results as the pyramidal cell population dominates the signal.

The reduction in the number of cells showing impaired activity but an increase in activity of individual neurons was perplexing, but may be explicable based on the complex pharmacological effects of muscarinic blockade. The reduction in cellular activity would be predicted from the inhibitory effect of muscarinic receptors on at least two types of potassium channels, especially the “M current” carried by Kv7 subtype channels inhibited by muscarinic receptors that are expressed at high density at the axon hillock, a specialised area of neurons modulating cell firing (Shah et al., 2008). ACh normally inhibits this current and increases activity, so scopolamine would be expected to reduce overall neural excitability. There are several other mechanisms by which muscarinic receptors increase neuronal activity (see Dannenberg et al., 2017), which would again likely produce reduced excitation with inhibition. On the other hand, Widman and McMahon (2018) have recently demonstrated in rat CA1 neurons *in vitro* reduced inhibitory input onto pyramidal cells and increased synaptically driven excitability measured at the single-cell and population levels which may be the mechanism for the increased activity seen. Sharp waves and ripples (SWRs) are hippocampal oscillatory patterns, which are believed to be crucial for memory consolidation, retrieval and planning (Girardeau et al., 2009; Carr et al., 2011). Although these oscillations are usually observed during animal’s behavioural immobility and slow-wave sleep (Buzsaki et al.,1992), they are also present during track or maze running period (Buzsaki, 2015; Cowen et al., 2020; see Joo and Frank 2018 for review). Some *in vivo* studies found that the reinforced hippocampal theta rhythm was characterized by the increased release of Ach (Parikh et al., 2007; Zhang et al., 2010), while inhibited cholinergic activity gave rise to SWRs (Buzsaki and Vanderwolf, 1983; Norimoto et al., 2012), and the suppression of SWRs could be rescued by muscarinic receptor antagonists (Ma et al., 2020). On one hand, blocking the muscarinic receptors increases the neural “M current”, which increases the firing potential threshold, leading to a reduction on the quantity of detected neurons. On the other hand, it also increases the probability of the appearance of SWRs, resulting in an increased average firing frequency.

Reduction of neuronal firing has also been seen *in vivo* with an mAChr antagonist (Brazhnik et al., 2003). If the neuron’s firing pattern became less stable, the *Ca*^2+^ event rate may increase, an extreme example was shown in Figure 11. In order to test this hypothesis, we studied the stability of neural ensemble. First, the spatial information content of the place cells reduced significantly after injecting scopolamine, which implied the fading location sensitivity of the neural ensemble, and clear differences could also be identified from the place field map. Besides, some researchers also found that the best fit line of the spikes recorded had a phase shift with respect to the corresponding local field potentials after injecting scopolamine in memory encoding and retrieval process (Douchamps et al., 2013; Newman et al., 2017), and this implied that the firing pattern of the neural ensemble changed after blocking mAChr, which consisted with our hypothesis to a certain extent. Interestingly, it seemed that the neural ensemble’s “spontaneous firing” was much stronger in the experiments when the animal run from left to right, possibly due to the clue painted on two ends (left: green colour clue; right: red colour clue) of the linear track were different, and the mice were more sensitive to green colour than red colour as described by Peirson et al. (2018). Finally, we quantified the stability of the neural system by calculating PVO between odd and even trials data within one session, the diagonalization feature disappeared after blocking mAChr. Besides, the value of mean PVO was relatively bigger with larger distance offset after blocking mAChr, this implied that the animal was more likely to confuse the locations that were far away from each other rather than adjacent locations in linear track.

**Figure 11.**
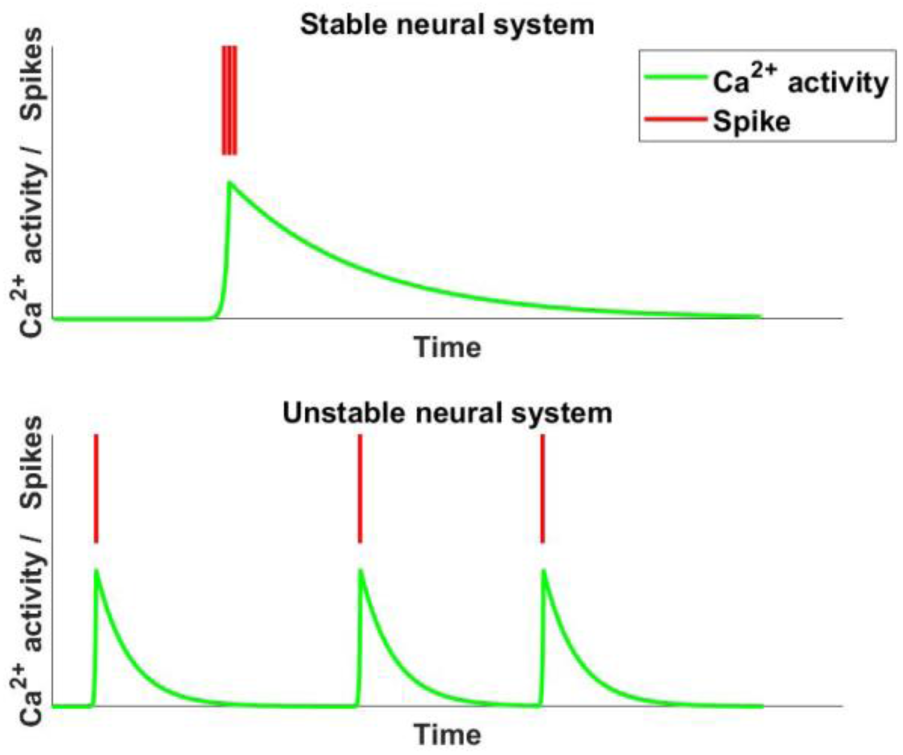
An extreme example showing that the *Ca*^2+^ event rate could increase with different neuron’s firing pattern. The calcium activity maybe submerged by continuous spike activity.

**Figure 12.**
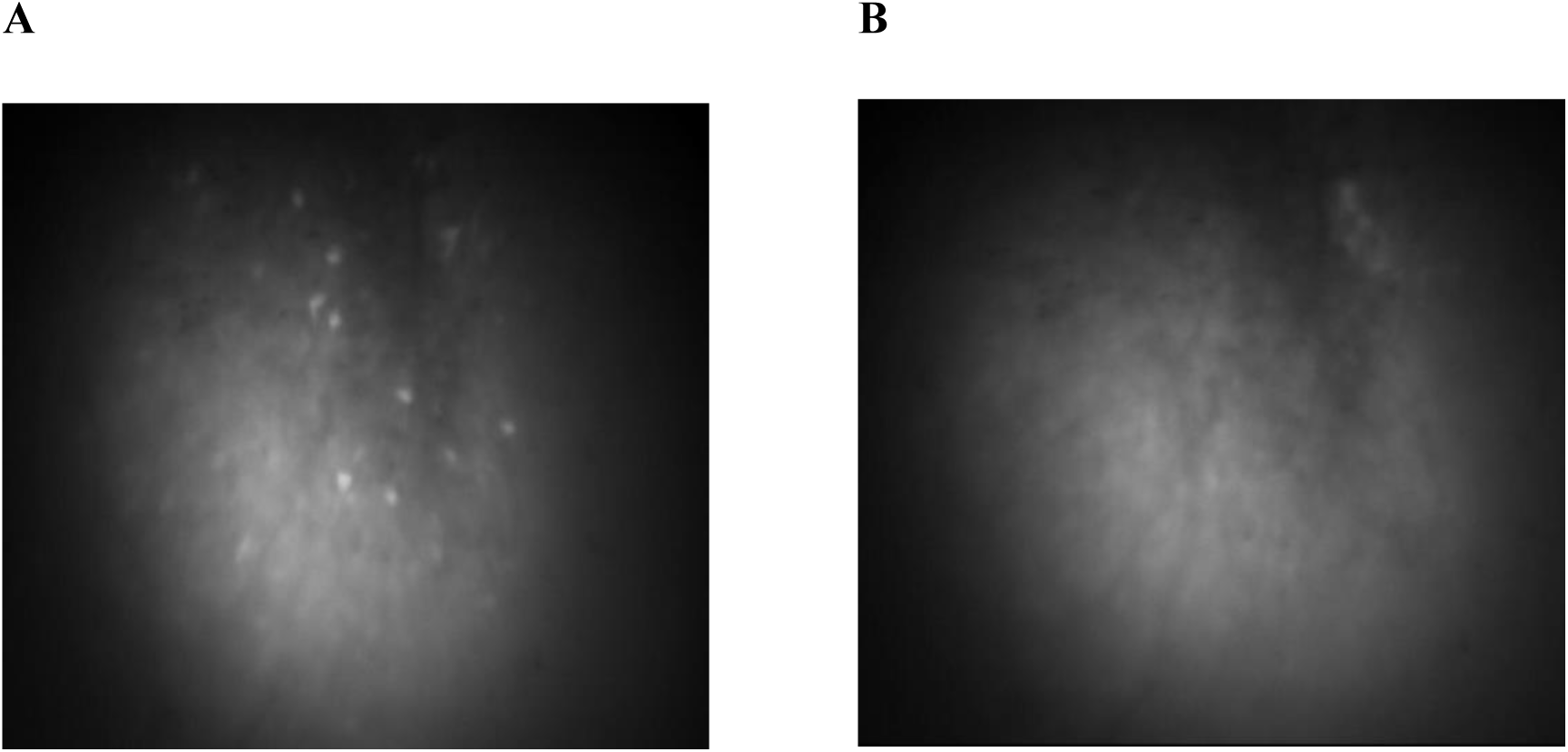
An example showing the *Ca*^2+^ activity before (A) and after (B) the administration of scopolamine. The fluorescent intensity decreased dramatically after injecting scopolamine.

The animal used in the experiments was trained to run back and forth in a linear track for multiple times and was given a long time period to acclimate the track before recording. A memory regarding the linear track should be formed and consolidated. This assumption could be verified from the results of saline group, the spatial information content of place cells and the mean PVO with respect to distance offset almost did not change too much compared with baseline. In order to make the most use of the data, we used a leave-one-out Bayesian classifier to estimate the animal’s position in the linear track. After the administration of scopolamine, the error rate increased dramatically, which consisted with the results of PVO map in some sense, and this implied that the mAChr antagonist had significant detrimental effects on memory retrieval. Cacucci et al. (2007) reported that the spatial tuning of place cells improved over time. Intriguingly, the saline group showed increased decoding accuracy, most likely due to the enhancement of the place fields with time.

The AAV virus we injected into hippocampal CA1 marked both pyramidal neurons and interneurons. Both pyramidal neurons and interneurons play important roles in animal’s spatial navigation, but with different mechanism of action (Gloveli, 2010), and the current miniscope recording system cannot distinguish them. As increased neural firing rate was observed after the administration of scopolamine, another possible reason may duo to the different effects of scopolamine on the firing pattern of pyramidal neurons and interneurons. The traditional method to distinguish the pyramidal neurons and interneurons by using intracranial electrodes depended on neuron’s mean firing rate (Fox and Ranck 1981; Ranck 1973), spike duration (Skaggs et al. 1996), and the autocorrelation function (Csicsvari et al. 1999). Sometimes, this statistics-based method cannot guarantee considerable accuracy, and simultaneous two-colour imaging (Aharoni et al., 2013) may be a possible better way to overcome this problem with the help of two carefully selected AAV virus combined with different fluorophores to mark the neurons separately. Both the new recording system and experiment design deserve further study.

In summary, our results suggest a close relationship between the blockade of mAChr and deficits in memory retrieval. The deteriorate memory decoding and increased *Ca*^2+^ event rate may derive from the decreased stability of the neural ensemble.

